# Acto-myosin network geometry defines centrosome position

**DOI:** 10.1101/2020.01.07.896969

**Authors:** Ana Joaquina Jimenez, Chiara de Pascalis, Gaelle Letort, Benoit Vianay, Robert D. Goldman, Michel Bornens, Matthieu Piel, Laurent Blanchoin, Manuel Théry

## Abstract

The centrosome is the main organizer of microtubules and as such, its position is a key determinant of polarized cell functions. As the name says, the default position of the centrosome is considered to be the cell geometrical center. However, the mechanism regulating centrosome positioning is still unclear and often confused with the mechanism regulating the position of the nucleus to which it is linked. Here we used enucleated cells plated on adhesive micropatterns to impose regular and precise geometrical conditions to centrosome-microtubule networks. Although frequently observed there, the equilibrium position of the centrosome is not systematically at the cell geometrical center and can be close to cell edge. Centrosome positioning appears to respond accurately to the architecture and anisotropy of the actin network, which constitutes, rather than cell shape, the actual spatial boundary conditions the microtubule network is sensitive to. We found that the contraction of the actin network defines a peripheral margin, in which microtubules appeared bent by compressive forces. The disassembly of the actin network away from the cell edges defines an inner zone where actin bundles were absent and microtubules were more radially organized. The production of dynein-based forces on microtubules places the centrosome at the center of this inner zone. Cell adhesion pattern and contractile forces define the shape and position of the inner zone in which the centrosome-microtubule network is centered.

## Introduction

The centrosome position is intimately associated to polarized cell functions such as adsorption and secretion, motility and mitosis [1–3]. Its position is characteristic and indicative of polarized cell functions [2,4]. It is found at the cell center in proliferating cells in culture [5] while it presents a somewhat peripheral position in differentiated cells in tissues, where it loses part or all of its microtubule organizing center (MTOC) functions [6]. In epithelial cells for example, the centrosome is found at the apical border of the cell and it nucleates few or no microtubules [7]. In ciliated cells, the centrosome role is more markedly changed as it turns itself into a basal body, responsible for the exclusive nucleation of cilia microtubules [6,8]. During several cellular events essential to development, and organism homeostasis, the centrosome position undergoes a shift from the center to periphery of the cell, notably during ciliogenesis [9], immune synapse formation [10] or Epithelial-to-Mesenchymal transition [11]. However the mechanisms that regulate the stability of central and peripheral states and those that allow a rapid switch between two states have not yet been fully understood.

Previous *in vivo*, *in vitro* and *in silico* studies suggest that centrosome position is the outcome of a balance of pulling and pushing forces applied on microtubules and transmitted to the centrosome [12–18]. Overexpression of dynein heavy chains, injection of dynein blocking antibodies, *in vitro* studies, and computational simulations suggest cortical and cytoplasmic dyneins play a role in the production of pulling forces [12,14,15,19,20]. Besides, microtubule polymerization against spatial boundaries have been shown to be responsible for the production of pushing forces [14,15,21,22]. Actomyosin contractility also modulates the forces applied on microtubules and can thereby influence centrosome position [23] although this parameter has remained controversial. The simple inhibition of actomyosin contraction had no visible effect on centrosome position in isolated cells or in monolayers [12,24], however it was found capable to counteract the asymmeric stimulation due local microtubule disassembly [12].

In non-differentiated cells, and notably in cells proliferating in culture, the force balance is believed to set the centrosome position at the cell geometrical center, also called center of mass or centroid [24–26]. Microtubule-based forces in an *in vitro* reconstituted system also position the MTOC at the centroid of their confinement area [14,15,27–29]. However, the centrosome has been observed at the cell geometrical center in relatively isotropic boundary conditions (e.g. non polarized cells) but can be off-centered in the front or in the back of migrating cells [30,31], or toward intercellular junction [11,32]. As a result, there is no generic definition of the centrosome position and the key parameters involved in the regulation of this positioning are still unclear.

One limitation for the comprehension of the forces exerted on the centrosome that ensure its location inside the cell is that the centrosome positioning is hardly distinguishable from nucleus positioning. It has been a considerable limitation for the study of centrosome positioning in anisotropic conditions such as in migrating cells [33,34]. Both the nucleus and the centrosome have their own self-centering properties [26,34–36] see (for Reviews [37,38]). But the physical links that connect them hinder their respective contributions in regards to their final position [38–45]. In addition, the nucleus also constitutes a dead volume microtubules don’t have access to, which biases the spatial distribution of microtubules and their associated forces. Furthermore, centrosomal microtubules push and pull on the nuclear envelop [38,46] adding more complexity to the force balance in the centrosome-microtubule network. For these reasons, enucleated cells –here referred as cytoplasts – offered an interesting possibility to untangle the geometrical and molecular cues that specifically control centrosome position. Plating them on on adhesive micropatterns revealed that centrosome self-positions at the geometrical center of the cytoplasts suggesting that its off-centering in cells is due to microtubule interaction with the nucleus[26,34]. However, the centrosome often detaches from the nucleus when moving to the cell periphery during the migration of neuroblasts [47] or epithelium formation [48] for example. This might indicate that the centrosome-microtubule network could be empowered of active off-centering properties independently of the nucleus, although this has not yet been demonstrated.

Here, we show that actin contractile network plays an important role in the confinement of the microtubule networks while the positioning of the centrosome at the center of this actin-based boundary is achieved by dynein-based forces on microtubules.

## Results

The major intermediate filament in fibroblasts, Vimentin, form a dense network around the nucleus [49]. Because of the elasticity and low dynamics of intermediate filaments, they recovered their initial shape after the enucleation step and formed a circular network with which microtubules kept interacting (Figure S1). To avoid any geometrical bias due to these structures interaction we worked with vimentin KO mouse embryonic fibroblast [50,51], in which centrosome position was similar to WT cells (Figure S1).

Cells were first plated on 2000μm2 disk-shaped micropattern, in order to maximise their spreading and the available space for centrosome positioning in 2D. Centrosomes were found to precisely position at the cell geometrical center: 84% were found in a 5μm wide region at the center of the disc (Figure 1A). Cytoplast, i.e. enucleated cells, were produced by centrifugation of attached cells on ECM-coated plastic slides [52]. They were then detached and plated on the same disk-shaped micropattern. Centrosomes displayed remarkably similar centering efficiency: 83% were found in the same central region (Figure 1A, Figure S1).

**Figure 1.**
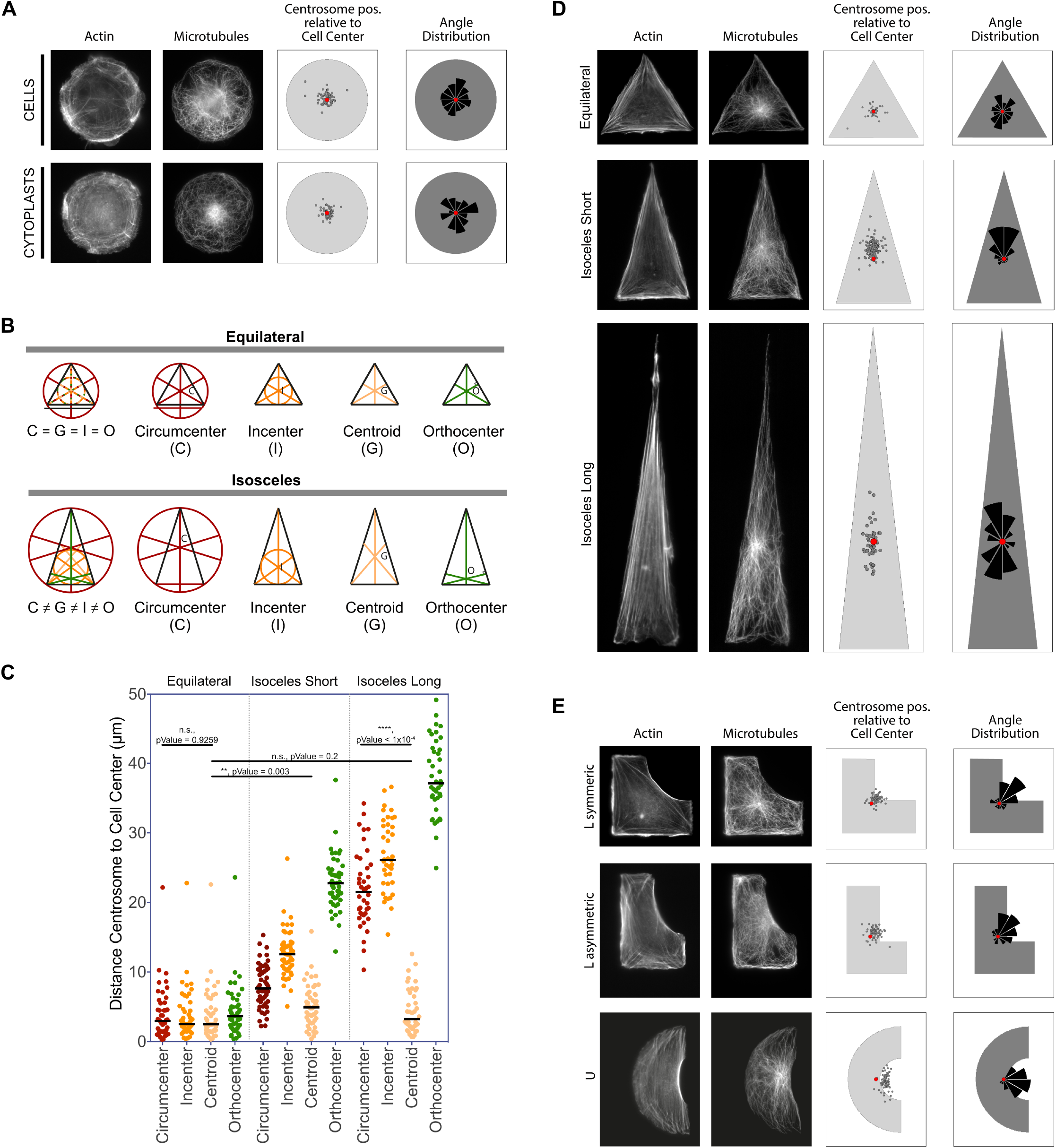
The centrosome located very close to the geometrical center of the cell. **A.** MEF KO Vimentin cells or cytoplasts plated on 2000μm2 disks for 4 hours, actin was stained with phalloidine-A555 and microtubules stained with rat anti-tubulin antibodies. **B.** Description of the triangle centers used for calculations of distances to the centrosome. **C.** Distance between the centrosome and the different triangle centers for three types of triangles, of 2000μm2: equilateral, isosceles 7/4 ratio (Short), isosceles 9/2 ratio (Long) reveals that the centrosome remains closer to the cell geometrical center than to the other centers especially when the aspect-ratio of the triangle increases. **D.** Actin and microtubule networks in the three kinds of triangles were stained as in **A**. The distribution of centrosome position relative to cell geometrical center shows a shift towards the cell apex in Isosceles “Short” and towards negative curvatures of the membrane in Ls and half-Thoruses. Red dots represent the cell geometrical center and grey dots represent the centrosomes.

In order to investigate centrosome positioning in anisotropic conditions, we plated cytoplasts on triangles. We chose triangles of similar area but different height to bases ratio: equilateral, short isoceles (ratio 7/4) and isosceles (ratio 9/2). Thousands of different geometrical centers have been described in triangles [53]. We chose some that are all are positioned along the midline of isoceles triangles, and could be considered interesting because their definition presumes particular interactions with the sides or the vertexes of the triangle. Those are the circumcenter (in close relationship with the triangle sides), the incenter (in close relationship with the triangle vertexes), the geometrical center (which reflects the whole area of the triangle) and the orthocenter. The distances between these centers increase with the height to base ratio of the triangle (Figure 1B), reflecting the variations of the contributions of their definition parameters (distance to vertex, distance to sides, distance to the middle of sides). Cytoplasts shapes were fitted with a triangular contour in order to measure the position of the various centers and compared them to the centrosome position. As the aspect ratio increased, the centrosome separated from all centers but the cell geometrical center with which it remained always in close proximity (Figure 1C). This confirmed the pre-existing definition of centrosome position in the field and suggested, by the geometrical definition of the geometrical center, that the whole spreading area of the cytoplast was essential to define the position of the centrosome. In order to investigate whether the centrosome would remain at the geometrical center of more complex shapes, we plated the cytoplasts on an exotic series of micropattern shapes and confirmed the robustness of centrosome positioning at the cell geometrical center (Figure S2).

However a small shift of the centrosome with respect to the cell geometrical was observed in the “short isosceles triangles” (Figure 1D, middle row). Interestingly, in these conditions, the actin network architecture was not homothetic compared to the cell contour (Figure 1D). The network width was larger along the triangle base and bundle arrangements at the small and large apices were different. These results suggested that centrosome position can be shifted from the cell geometrical center and this could be due to higher asymmetry in the actin network.

Previous works showed that the presence negative curvature (concave edges) in cell attachment areas induces anterograde flow of actin from the areas of positive curvature (convexe edges) while static stress fibers are generated at the areas of negative curvature [54,55]. We thus plated cytoplasts on shapes made of combination of concave and convex edges to induce a strong asymmetry in the actin network architecture. In “L” shapes or “half-thorus” shapes, actin network displayed a marked asymmetry as expected and the centrosome was significantly shifted from the cell geometrical center, away from the retrograde flow and towards the stress fibers (Figure 1E). We were able to conclude that the centrosome position depends on the cell adhesion pattern and the architecture of the actin network, regardless of the nucleus.

Interestingly, a central zone in the cell is devoided of actin bundles: we called it the Actin Inner Zone (AIZ). Using image denoising and edge detection we semi-automatically detected the contour of this zone and its geometrical center: the Actin Inner Center (AIC) (Figure 2A). We were able to conclude that the centrosome is closer, or equally distant, to the AIC than to the cell geometrical center in the geometrical shapes we tested (Figure 2B). Therefore, the AIC appeared to be a better descriptor of centrosome positioning than the cell geometrical center. We thus further investigated whether microtubules engaged specific and distinct interaction in and out of the AIZ.

**Figure 2.**
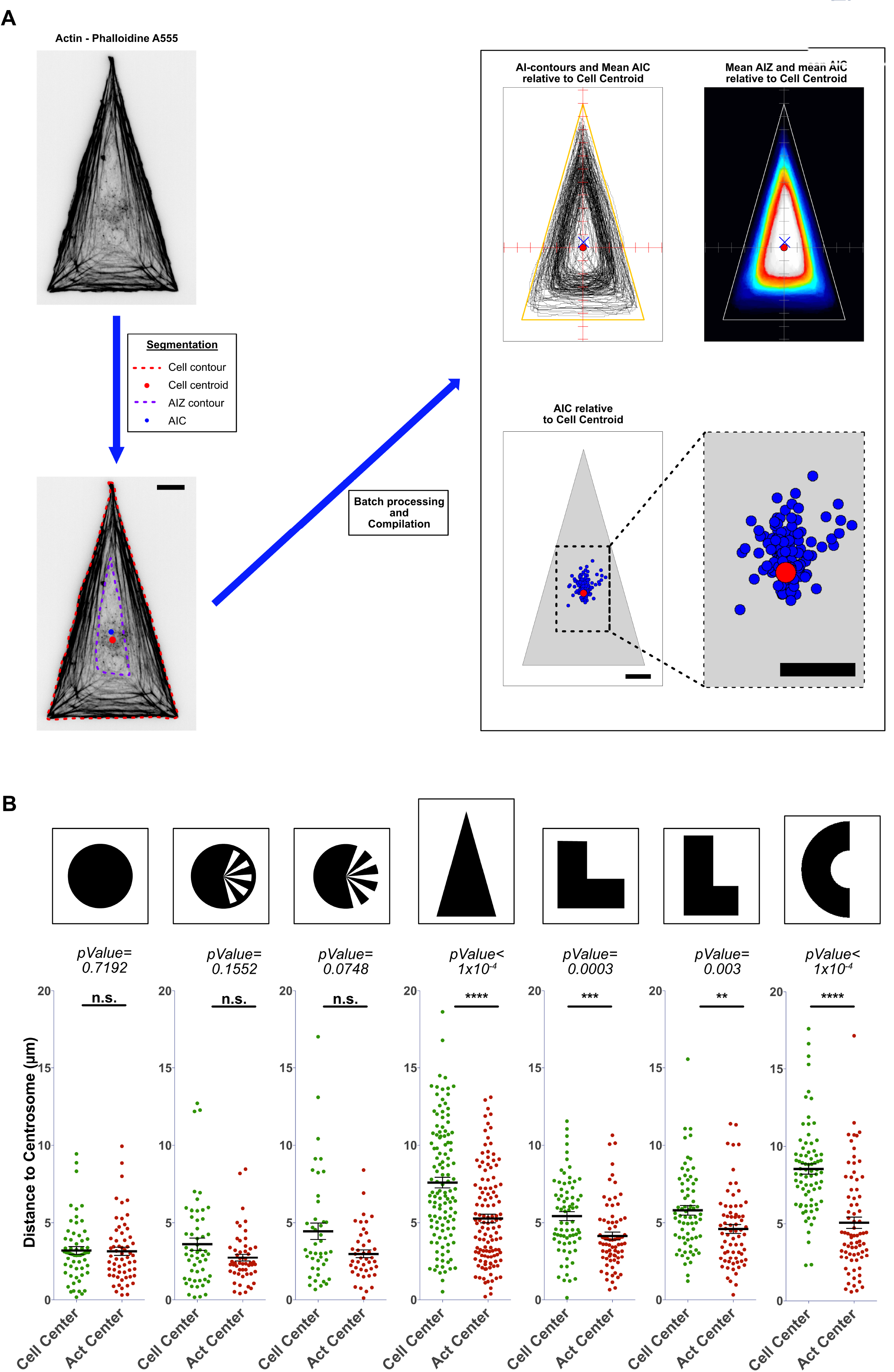
The centrosome located very close to the geometrical center of the cell. **A.** MEF KO Vimentin cytoplasts were plated on isosceles “Short” triangles, fixed and stained with Phalloidine-A555. The scheme shows the analysis performed to study the AIZ and AIC. The red dot represents the geometrical center of the cell and the blue cross represents an averaged AIC relative to cell center. **B.** The distance of the centrosome to the cell geometrical center was compared to the distance of the centrosome to the AIC for several shapes. The most significative are presented. Even if for some shapes the difference is not statistically different, all shapes present a distance centrosome-to-AIC equal or shorter than the centrosome-to-cell center distance.

To quantify the influence of regions of actin retrograde flow and regions devoided of it on microtubule organisation and the centrosome positioning, we developed a new image analysis tool “D-FiNS” (for Democratic Filament Network Scanner) soon available on gitHub. The steps of this analysis are described in Figure 3A. D-FiNS allowed us to detect and segment, in batch, actin bundles and microtubules in epifluorescence stack images and measure local microtubule orientation (Figure 3A, Figure S3). These orientations were clustered in two categories: radial (<45°) and tangential (>45°). The averaged local orientation was further defined by the “Orientation Ratio” (OR) as the ratio of pixels with radial over tangential orientation. The OR was established for the entire cell or along radial linescans. We then compared microtubule orientation in the AIZ and out of the AIZ (Figure 3B). Clear differences revealed that microtubules were radially oriented in the inner part of the cell when devoided of actin bundles and more tangentially oriented along with actin bundles, at the cell periphery (Figure 3C, Figure S4). From these results we concluded that the architecture of the actin network acts locally on the shape and orientation of microtubules.

**Figure 3.**
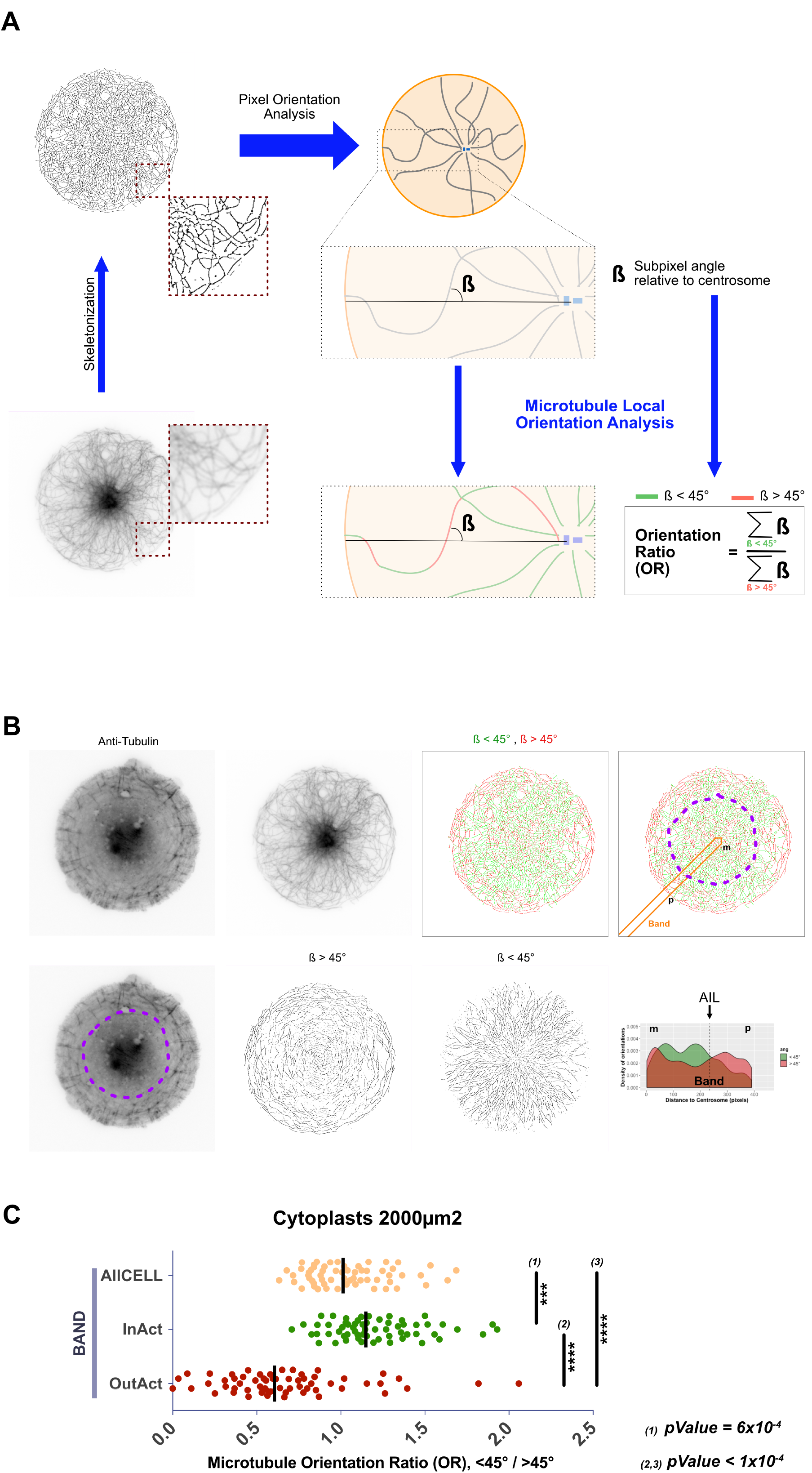
Microtubule network analysis and cross-analysis with AIZ. Cytoplasts were plated on 2000μm2 disks and stained for Actin and Microtubules like in Figure 1. **A.** Overview of the microtubule network analysis using D-FiNS (shortly available for the community). The Orientation Ratio or “OR” is the ratio of the number of non-null pixels with an orientation <45°C and the number of non-null pixels with an orientation >45°C. **B.** Overview of cross-analysis: microtubule network analysis along a radial band and inside and outside the AIZ. AIL stands for Actin Inner Limit. The AIL is found at the boundary of two zones defined by a clear majority of radial or tangential microtubules respectively, showing the strong correlation between actin and microtubule network architecture. **C.** Results of cross-analysis plotting the “OR”. The microtubules within AIZ are preferentially oriented radially and the ones outside the AIZ are preferentially oriented tangentially.

The role of the size of the AIZ was further confirmed by experiments on various sizes of discs ranging from 500 to 3000μm2. Surprisingly, we found that the width of the distribution of centrosome positions was independent from the size of the disc (Figure S5A, B, C). Parallel to this, we also found that the size of the averaged AIZ was relatively constant in cytoplasts with different sizes (Figure S5D). This suggests that the centrosomes and the microtubules were more sensitive to the AIZ than to the cell periphery. To further confirm this model and test the impact of the AIZ on centrosome position, we modulated the shape and position of the actin bundles and the AIZ. We further worked with short isoceles triangles as those shapes were shown to shift centrosome position away from the geometrical center (Figure 1D) while ensuring a more efficient cytoplast spreading than on L or half-thorus shapes. Disassembling actin perturbs too much the spreading of cytoplasts. Arp2/3 and Formin inhibition resulted in perturbation of cell shape and partial disruption of microtubule network respectively, precluding their use. Interestingly, inhibiting the Rho kinase ROCK with Y27632, inhibited the assembly and retrograde flow of actin bundles resulting in an homothetic network of thin and loose actin bundles along all cell edges (Figure 4A). In particular, the AIZ was enlarged and the width of the network along the short edge was reduced compared to control conditions and similar to the width along the two other edges (Figure 4B). As a consequence, the center of the AIZ, which was shifted from the cell geometrical center in control conditions, was moved back to this center in response to ROCK inhibition (Figure 4C). The centrosome position responded to ROCK inhibition as well and like the AIC it was shifted back to the cell geometrical center, (Figure 4D,E). We were able to conclude that the actomyosin contraction defines the shape and position of the AIZ, which acts as spatial boundary conditions to the microtubule network and thereby impacts centrosome position. The following step was to investigate how the centrosome could position at the center of the AIZ.

**Figure 4.**
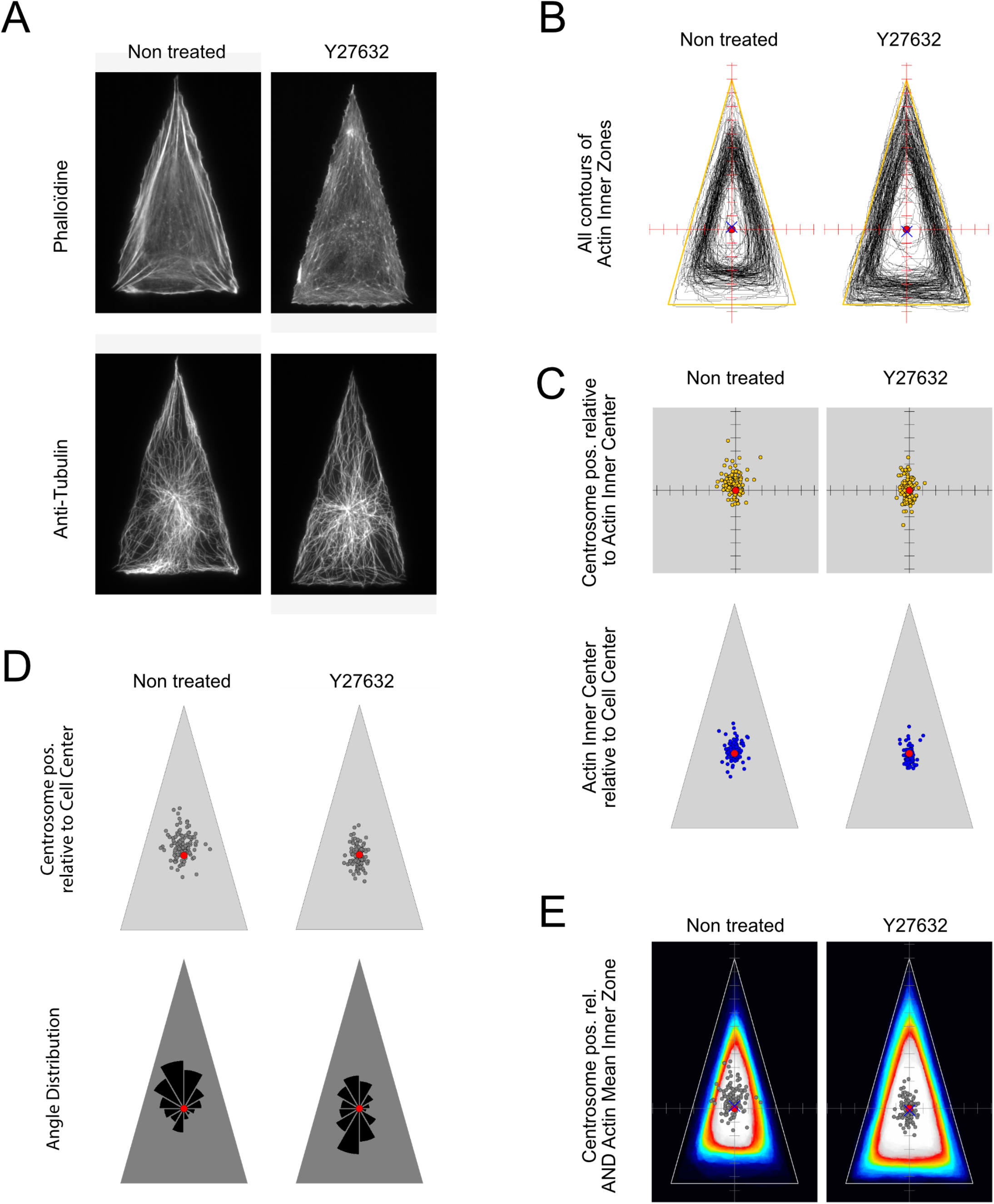
Role of contractility in AIZ/AIC and centrosome positioning. Cytoplasts plated on 2000μm2 Isosceles “Short” triangles were treated for 2 hours with Y27632 at 20μM. Analysis was performed as described before. **A.** Microtubule and actin staining as in Figure 1. One representative example is given for each condition. Stress fibers are observable in the control condition, with higher contractility coming from the basis of the triangle. In cells treated with Y27632, actin presents a rather branched meshwork architecture that extends from the edges of the cell in a homogenous/homothetic manner towards the inside. **B.** Plots of all the contours of AIZs relative to the cell center. The contours are larger and homothetic with the triangle edges in Y27632 conditions. **C.** The AIC is better centered in Y27632 conditions. The position of the centrosome relative to the AIC is similar in both conditions. **D.** Centrosome is relatively well centered (close to the cell geometrical center) but a small shift is observable towards the triangle apex in control cells. Its centering is improved with the Y27632 treatment. **E.** The averaged AIZ is wide, triangle-shaped and centered in Y27632 conditions, while it presents a less clean triangle shape in control conditions where it is slightly shifted towards the cell apex and smaller, reflecting higher contractility. In both cases, the centrosome distribution is found inside the common area of most AIZs, but it is more dispersed in control conditions. Red dot represents cell geometrical center or the AIC as indicated by the graph title. In E, the red dot represents the cell geometrical center and the blue cross represents the averaged AIC relative to cell center. The grey dots are the centrosomes.

The centrosome position depends on the mechanical forces exerted on the microtubule network [12,13,17–19,22,56]. Microtubule disassembly is known to impact the centrosome position [24,34]. We tested the impact of microtubule disassembly on centrosome position relative to the overall cell shape and relative to the AIZ. Cytoplast were plated on ice (2hours) and treated with 10μM nocodazole to induce a complete disassembly of microtubules (Figure 5A, left). This treatment induced a significant dispersion of centrosome positions (Figure 5B, left). Importantly, and in accordance with previous studies, microtubule depletion also increased cell contractility [57] and induced the formation of large actomyosin bundles (Figure 5A, left). As a consequence, the AIZ was severely deformed and shifted asymmetrically with respect to the overall cell shape (Figure 5, leftC). In such asymmetric networks the AIC was moved quite far away from the cell geometrical center (Figure 5D, left), an effect that could have had an additive contribution to centrosome dispersion.

**Figure 5.**
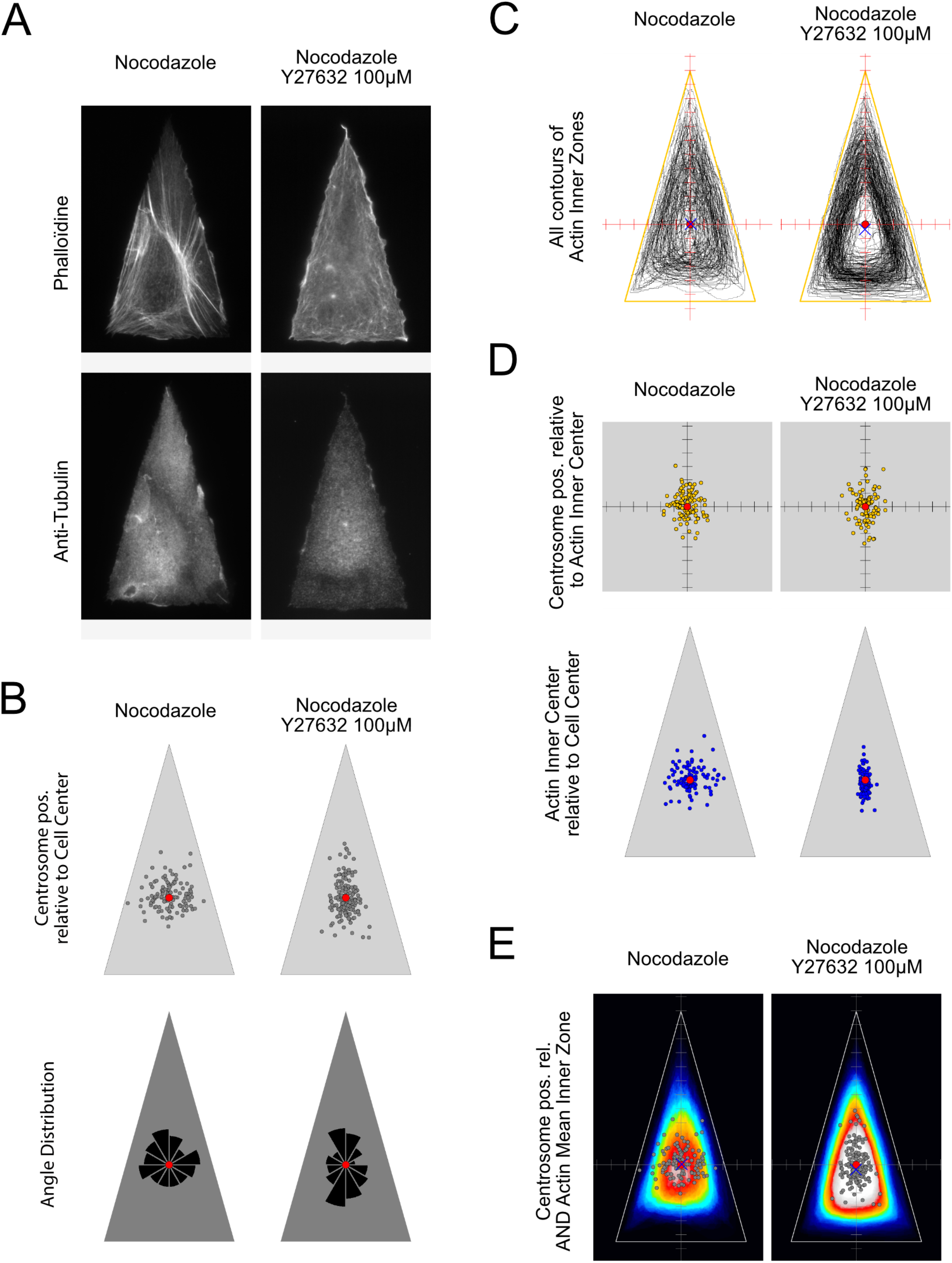
Effect of microtubule depletion on AIZ/AIC and centrosome positioning. Cytoplasts plated on 2000μm2 Isosceles “Short” triangles were treated with Ice-Nocodazole 10μM or Ice-Nocodazole 10μM/Y27632 100μM in order to uncouple the effect of contractility and the one of microtubules depletion. Control conditions were systematically performed but are not systematically shown to avoid message redundancy and overcharged figures. **A.** Representative images of cytoplasts from both conditions. In the absence of microtubules, actin contractility is highly increased. In cells depleted for microtubules and treated with Y27632 at 20μM, contractility was still abnormally high (not shown). A higher concentration of Y27632 (100-200μM) was then used. As observed in the example cell, this concentration was sufficient to inhibit contractility in the absence of microtubules. **B.** In the absence of microtubules the centrosome distribution is dispersed around cell geometrical center. The additional treatment with Y27632 maintained the dispersion but with a different anisotropy suggesting that part of the effect upon removal of microtubules is due to contractility. **C.** In the absence of microtubules, AIZ contours are more off centered than in control conditions, revealing the presence of anisotropic contractility, which could be responsible for the off-centering of the centrosome. This is also revealed by the increased dispersion of AICs (panel **D**). **D.** Both conditions show increased dispersion of the centrosome positioning relative to the AIC and compared with control condition and condition with no contractility (Figure 4C), but less dispersion than for its position relative to cell geometrical center. This suggests that, in the absence of microtubules, the centrosome lacks centering mechanisms within the AIZ. **E.** Overlapping decrease is revealed by averaged AIZs in Nocodazole conditions while the addition of Y27632 rescues this effect. Interestingly, centrosome dispersion forms a triangle shaped region, which fits in the overlapping region of all AIZs. Red dot represents cell geometrical center or the AIC as indicated by the graph title. In E, the red dot represents the cell geometrical center and the blue cross represents the averaged relative AIC. The grey dots are the centrosomes.

Interestingly the effect of microtubule disassembly on centrosome positioning on disks was also consistent with a role of the increased contractility and AIZ distortion. Microtubule disassembly did not completely randomized centrosome position even after more than 20h of treatment (figure S6A, B). The distribution was only twice larger. In parallel the AIZ was reduced by the increased contraction up to a size that appeared consistent with the limited dispersion of centrosomes (Figure S6C). Therefore, and importantly, both experiments suggested that the well-known centrosome mispositioning in response to microtubule disassembly resulted not only from the absence of microtubule but also from the deformation of actin-based spatial boundaries.

In order to ungroup those two effects and test specifically the impact of microtubule disassembly without changing AIZ conformation, we set up conditions to disrupt both MTs and compensate for actin contractility by using ice-cold/nocodazole treatment combined with high doses of Y27632 (100 or 200μM) (Figure 5A, right). In these conditions, the AIZ was similar to the conditions described previously in response to Y27632 alone (at 20μM): it formed a regular, homothetic peripheral band along the cell edges (Figure 5C, right). In those conditions, centrosomes dispersion appeared limited within the AIZ (Figure 5E, right) and not randomly distributed in the entire cell. In conclusion, in the absence of microtubules, the centrosomes cannot diffuse all over the cell and remain confined in the AIZ. We then further investigated how microtubules direct centrosome position at the center of the AIZ.

Dynein is involved in centrosome positioning in eggs [17,56], embryos [58] and mamalian cells [12,19]. By acting at the cell cortex or in the entire cytoplasm, dyneins are thought to position the centrosome at the geometrical center of the cell. We tested ciliobrevin D and dynarrestin but found no clear effect on the dispersion of the Golgi apparatus. So we chose to inhibit dyneins activity by expressing a dominant negative form of the dynactin subunit p150glued (p150-DN) [59], which is involved in connecting dyneins to cargo and provide a load to motor activity. p150-DN was transfected in cells 48 hours before enucleation. In these conditions the centrosomes were dispersed but again, did appeared to become randomly throughout the entire cell. Their distribution was still anisotropically biased in regards to cell geometrical center (Figure 6A). This effect was consistent with the maintenance of a shifted AIZ similar to control conditions (Figure 6B) in which the centrosome appeared dispersed (Figure 6C).

**Figure 6.**
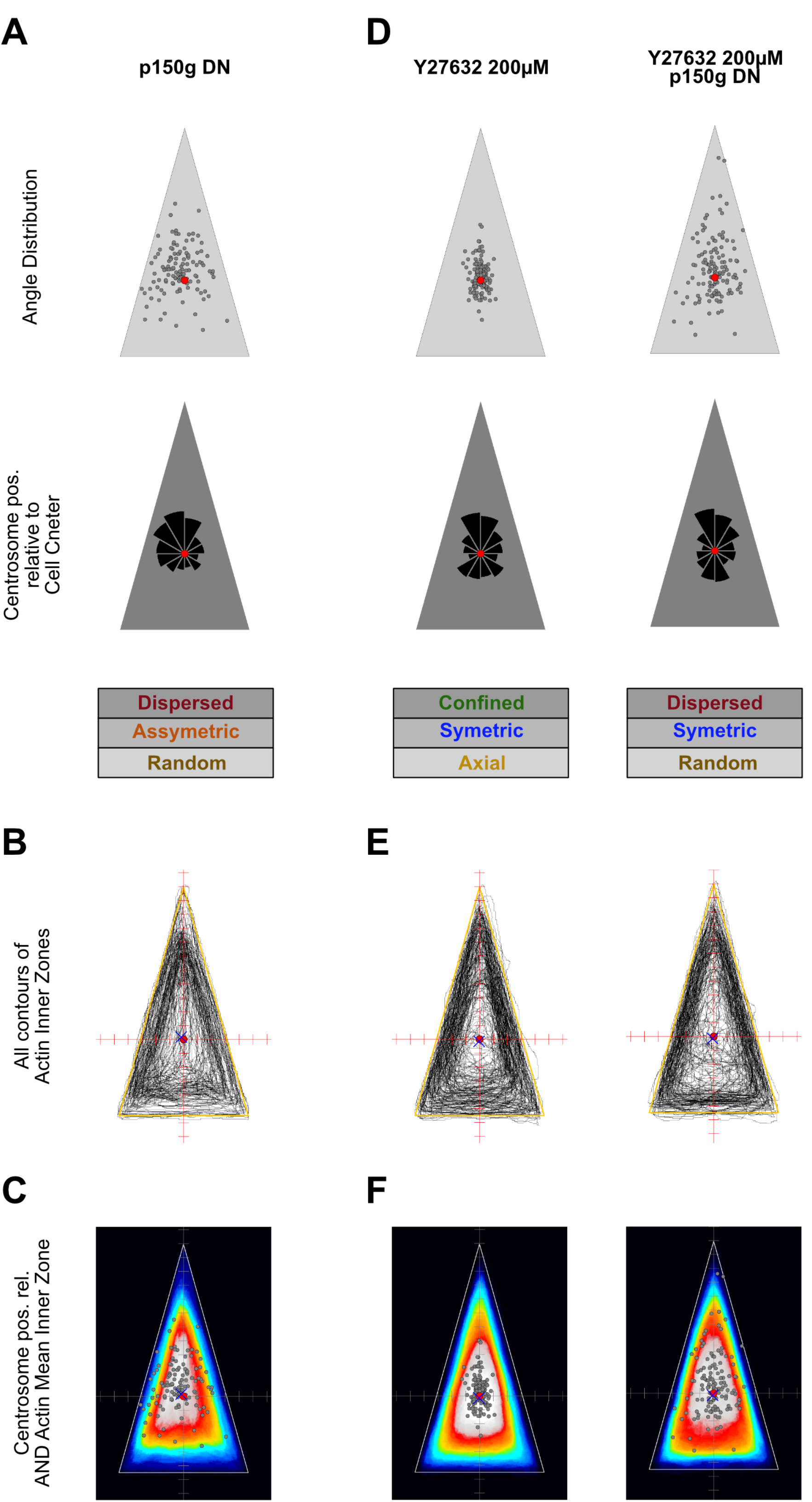
Role of dyneins and microtubule depletion in AIZ/AIC and centrosome positioning. Cells electroporated with p150-DN 48 hours before were enucleated. Cytoplasts were plated on 2000μm2 Isosceles “Short”. **A, D.** The centrosomes are dispersed when p150-DN is expressed, in the presence or absence of contractility, all over the cell. **B, C,E,F.** AIZ contours are off-centered similarly to Figure 5C, while in the presence of Y27632 at 200μM, they follow mostly cell edges. Centrosome distribution is not fully confined in AIZs overlapping region. Therefore, dynein inhibition before cytoplasm spreading on patterns affects centrosome inclusion into the AIZ. Red dot represents cell geometrical center or the AIC as indicated by the graph title. In E, the red dot represents the cell geometrical center and the blue cross represents the averaged relative AIC. The grey dots are the centrosomes.

In order to further test whether dynein helps to find the center of the AIZ, we tested the role of dynein activity in conditions imposing another shape and position to the AIZ, i.e. in the absence of cell contractility (Figure 6D-F, right panels). As described previously (Figure 4B), in these conditions, and in contrast with cells with normal levels of contractility, the actin network forms a regular band along cell edges, and thus the center of the AIZ is moved back to the cell geometrical center (Figure 6E). In Y27632-treated cells centrosome positions were distributed around the cell geometrical center regardless of the activity of dyneins. However they were concentrated around it when dynein were actively pulling on microtubule (Figure 6D, left) whereas they were dispersed over the entire AIZ when they were inactivated (Figure 6D, right). Again, dispersed centrosomes remained confined in the AIZ (Figure 6F). These results showed that dyneins activity is involved in the positioning of centrosomes at the center of the AIZ.

## Discussion

### Centrosome, nucleus and the cell geometrical center

The mechanism which specifically regulates the positioning of the centrosome in mammalian cells has long been confused with the role of the nucleus [53], and notably the role of actomyosin network acting on the nucleus [19]. The consensus was that the centrosome positions at the cell geometrical center, either autonomously [21,30] or in association with the nucleus [19]. Here we studied the centrosome separately from nucleus and showed that the centrosome position is defined by the architecture of the actin network. It is located at the center of the actin network; more precisely, it is positioned at the geometrical center of the inner space that is devoided of actin bundles. This position can correspond or not to the cell geometrical center, depending on the anisotropy of the actin network.

### Microtubule and actin

Microtubules interact with actin via specific crosslinkers or non-specific steric interactions [61]. In particular dense and growing actin network can apply pushing forces on MTs [62,63]. Here we found that the actomyosin network constitutes the actual spatial boundary conditions that the microtubules are sensitive to. Microtubules appeared bent within the actomyosin network, and straight in the central part devoid of actin. Disrupting or moving the geometry of the network via the pattern of adhesion moved this boundary and changed centrosome position accordingly.

### Dyneins

Dynein have long been known to apply pulling force on MT [64] and to be thus involved in MTOC positioning [12,14,15,19,20,65]. By acting at the cell cortex, or throughout the cytoplasm, dyneins are thought to position the MTOC at the cell geometrical center [66]. Here we found that dyneins are involved in centrosome positioning at the center of the inner cell space that is devoid of actin bundles. The radial shape of microtubules suggests that dyneins put them under tension in this region. Dynein activity requires dynactin to be coupled to a cargo or any other substrate supporting the force load. It is not clear whether these dyneins act throughout this region via cytoplasmic forces [67] or only at the interface between the actomyosin network and the microtubule network, where they have been proposed to act as acoupling device that transmits contractile forces from the actomyosin network onto microtubules [58].

### Implications for centrosome positioning in differentiated/polarized cells

By using enucleated cells to distinguish the centrosome from nucleus positioning this work reveals that centrosome is not at the cell geometrical center but at the center of the actin network. The actin-centering forces are potent and significant since highly asymmetric actin networks, such as those developed on half-thorus shaped micropatterns, can force the centrosome to come into contact with cell periphery. These results revealed a novel positioning mechanism, which shines some light on numerous conflicting examples of centrosome positioning in migrating or polarized cells (cf introduction). They show that the organization of actin filaments, driven by myosin-based contraction, are the actual boundary conditions that influence microtubule organization and direct centrosome position.

## Experimental Procedures

### Cell culture, cell lines, plasmids and transfection and drug treatment

MEF WT and KO for Vimentin cell lines (received from Robert Goldman) were cultured in Dulbecco’s Modified Eagle Medium (31966, Gibco) supplemented with 10% FBS (50900, Biowest) and 1% antibiotic-antimycotic (15240-062, Gibco). RPE1 cells were cultured in Dulbecco’s Modified Eagle Medium Nutrient Mixture F-12 (31331-093, Gibco) supplemented with 10% FBS and 1% antibiotic-antimycotic. MEF KO Vimentin EGFP-Centrin 1 were made by transient transfection of pEGFP-C1-Centrin1 (kindly provided by James Sillibourne) with lipofectamine LTX (15338100, Invitrogen) in Opti-MEM (11058, Gibco) according to the procedure described by the manufacturer. Selection was performed with G418 at 0.5mg/ml and sorted by FACS twice with one month interval. They were posteriorly cultured with 0.2mg/ml. For p150 inhibition assay on cytoplasts, cells were electroporated with NEPA21 electroporator (from Nepa Gene) with the plasmid expressing GFP-p150-CC1 (214-548 aa of p150Glued) obtained from Mineko Kengaku (Kyoto University) and according to the manufacturer’s protocole for MEF cells. Cells were then sorted by FACS 24h after electroporation and plated directly on slides for enucleation and enucleated 48h after electroporation. Living cells were incubated and imaged at 37°C with 5% CO2 in a humidified environment.

### Enucleation

Cells were seeded the night before, 12h before enucleation on RINZL plastic micro-slides (71890-01, Delta Microscopies) precoated with Collagen I Rat Protein, Tail (A1048301, Gibco) at 12μg/ml and Fibronectin from bovine plasma (F1141, Sigma) at 1μg/ml for 1 hour. Cells were seeded to achieve a 90 % confluence by the time of the enucleation. Cells were put on 50ml tubes resistant to high-speed centrifugation (339652, Nunc) in complete medium with Cytochalasin D (C8273, Sigma) at 3μg/ml for 30min at 37°C, then centrifuged at 15’000 g for 1h at 37°C. Cytoplasts were then washed twice with pre-warmed DMEM then let them to rest for 30min at 37°C before detachment for seeding on micropatterns.

### Micropatterning and cell seeding

#### Polystyrene coating

20Ô20 Coverslips (1304369, Schott) were cleaned for 10min in acetone then for 10min in isopropanol in a bath sonicatorn then compressed-clean air dried under a laminar flow hood. They were coated with adhesion promoter Ti-Prime (MicroChemicals) using a spin-coater (WS-650m2-23NPPB, Laurell) at 3000 rpm for 30s. and baked on top heater for 2min at 120°C, followed 1% polystyrène (MW 260,000, 178891000, Acros Organic) solution in toluene (179418, Sigma) using a spin-coater at 1000rpm.

#### Plasma treatment and micropatterning

Polystyrene layer was oxidized by exposure to air-plasma (PE-30,Plasma Etch) at 30W, under vaccuum and with an air flow rate of 10 cc/minute, for 40 seconds to promote the attachment of Poly-l-Lysine-Polyethylenglycol/PLL-PEG (PLL(20)-g[3.5]-PEG(2), SurfaceSolutionS, Switzerland), which was performed in a solution of HEPES at 10mM, pH7.4, with a concentration of PLL-PEG at 0.1mg/ml for 30minutes at room temperature. PLL-PEG was removed and coverslips room air dried before putting them in tight contact with a chromed printed photomask (Toppan Photomask). Tight contact was maintained using a vacuum holder. The PLL-PEG layer was burned with deep UV (λ=190nm) through the non-chromed windows of the photomask, using UVO cleaner (Model No. 342A-220, Jelight), at a distance of 1cm from the UV lamp with a power of 6mW/cm2, for 4 min.

#### Cell seeding

Coverslips were washed once with distilled water then incubated with a solution of 40μg/ml Fibronectin (F1141, Sigma) in PBS (14190169, GIBCO) for 30min at room temperature. Coverslips were then washed, in a sterile 6-well dish with one coverslip per well and under the laminar flow hood, 3 times with 3ml sterile PBS, once with 3ml DMEM and once with 3ml DMEM-10%FBS-1%Antibiotic-Antimycotic (complete medium). Cells/cytoplasts were detached with TrypLE (12605036, Gibco), centrifuged and resuspended in complete medium at 100’000 cells/ml. Most medium was removed for each well containing a coverslip and 1ml of cell suspension was added. Cells were left for spreading for 1 hour at 37°C before washing-out non-attached cells with pre-warmed complete medium. Cells were incubated for at least one more hour at 37°C to promote correct spreading and polarisation, before further treatments.

### Drug treatment

Microtubules were removed by incubating cells in HBSS (14025092, Gibco) on ice and in a cold room at 4°C for 2 hours then warmed up to 37°C in complete medium with 10μM Nocodazole (M1404, Sigma) and incubated until fixation. Rock inhibition was achieved with Y27632 (Y0503, Sigma) at 20μM. Rock inhibition in the absence of microtubules was achieved with cold incubation as described above and warming up with complete medium with 10μM Nocodazole and Y27632 at 100 or 200μM as specified for 2h at least.

### Immunostaining

Cells plated on coverslips were fixed with Paraformaldehyde (15710, Euromedex), Glutaraldehyde (G5882, Sigma) or a mixture of both depending on the antibodies used. All fixation mixtures were done in Cytoskeleton-Sucrose buffer with 0.1% Triton X-100 (T8787, Sigma) with either 3% Paraformaldehyde, 3%Paraformaldehyde-0.025% Glutaraldehyde or 0.5% Glutaraldehyde. Fixation mixture was added to the cells for 10min at room temperature. Glutaraldehyde related autofluorescence was quenched with a solution of PBS and 1mg/ml sodium Borohydride for 10min at room temperature. Cells were then re-permeabilised with PBS-Triton 0.1% for 10min at room temperature, then in PBS with Bovine Serum Albumin/BSA (A2153, Sigma) at 1.5% for 10min. Antibodies were diluted in PBS-BSA 1.5% and both incubation with primary or secondary antibodies was made for 1 hour. Microtubules were stained with MCA77G from Abd serotec or with ab18251 from Abcam. Centrosome staining was performed with anti-gamma Tubulin (T6557,Sigma), anti-pericentrin (ab4448, Abcam) or anti-polyglutamylated Tubulin (A-R-H#04, TabIP platform, Institut Curie). Actin filaments were stained with Phalloidin-A555 (A34055, Life Technologies) or Phalloidin-A568 (A12380, Life Technologies) together with secondary antibodies. Staining with DAPI (D9542, Sigma) was performed systematically with secondary antibodies to control proper enucleation. Coverslips were mounted with Mowiol 4-88 (81381, Sigma).

#### Cytoskeleton – Sucrose buffer

A stock solution containing 10mM HEPES (H3375, Sigma) at pH 6.1, 138 mM KCl (P3911, Sigma), 3 mM MgCl2 (208337, Sigma) and 2 mM EGTA (E3889, Sigma) was made. Sucrose was added extemporaneously at 0.32M.

### Microscopy

Fixed and fluorescently labeled cells were imaged using an up-right epi-fluorescence microscope (Olympus up-right BX61) monitored by Metamorph. Samples were scanned for cell selection with dry objectives 10x or 20x using a Metamorph plugin developped by Céline Labouesse and Benoît Vianay. Cells were chosen so that they were well spread on sharp patterns and that they do not had a nucleus in the case of conditions with cytoplasts. Cells were imaged with a 100x NA 1.4 oil objective, with 0.5μm spacement between z planes in a range of 15 μm. When patterned cells did not fit in one camera field, overlapping images were taken for further stitching.

### Image analysis

#### Image analysis of very big cells

For patterned cells that could not fit in one camera field, ImageJ macros using Stitching plugin were used. Images were then processed the same than single images.

#### Centrosome positioning compared to cell centroid and nucleus centroid

This analysis was performed with homemade ImageJ suite of macros. The closest plane to the coverslip (cell bottom) was determined creating a band ROI on the actin image, as an expansion of a rough cell border determined by threshold filtering. This ROI was applied to the microtubule channel where the z-plane with the highest Standard Deviation within the band was chosen as cell bottom. Cell Top was determined using the standard deviation of the whole image. Firsts and lasts superfluous z-planes were that way removed to lighten calculations.

Threshold filtering in actin cell bottom plane was performed to determine cell edges and centroid was calculated. Similar method was used to determine nucleus edges and centroid in the case of conditions with nucleated cells. Scanning the Prominence parameter of the “Find Maxima” plugin from imageJ was performed to determine the prominence value where the number of found maxima was closest to one. Scanning for maxima within the region around this principal centriole was performed to find eventual extracentrioles. Cells with more than 4 centrioles were discarded. The centroid of the polygon defined after connecting all centrioles 3 by 3 into triangles and adding all areas was used for the calculation of the distances to the cell, and nucleus if applicable, centroids.

All steps contained a quick-scanning verification and assisted-correction module to make sure the analysis was correct for all cells.

#### Centrosome positioning compared triangle characteristic centres

In the case of triangle-patterned cytoplasts, the contour defined previously was smoothened by converting curve into a spline defined by a discrete number of close points. The curve defined by the distance of each point from the spline to the previously calculated centroid was smoothened by quadratic regression until the curve presents only 3 maxima, corresponding to the 3 triangle vertexes. The indexes of these three points were used to find the 3 corresponding points the contour-spline. The coordinates of theses 3 points were used to fit the contour of the cell to a triangle. Geometrical calculations were performed to determine the coordinates of 4 characteristic centres of that triangle (centroid, incentre, circumcenter and orthocentre). Distances from the centrosome to these centres were calculated.

#### Actin inner zone (AIZ) and actin inner centre (AIC)

Actin inner zone was determined semi-automatically on projected and denoised (rolling ball filtering) actin images. The coordinates centroid of the zone was determined and the distance to cell and centrosome centroids was calculated.

#### Dot plots and plots of AIZs

An angle correction was determined for all cells in a semi-automatic way. The coordinates of all centres were redressed according to the correction angle and relative coordinates to cell or actin centroid were calculated and plotted. Similar procedure was performed for the regions defining AIZs. Either all contours of AIZs were drawn, or one black 8-bit image was created for each cell and the AIZs was drawn and filled in white. A sum of all the images was made and a Royal LUT was applied.

#### Microtubule orientations

Microtubule stacks were skeletonized using a homemade Java plugin. A sum projection was made before a homemade orientation filter was applied to determine the angle made by each pixel of the skeletonized microtubule network. The calculation of a relative angle to the centrosome was performed. This angle corresponds to the angle made locally by a portion of microtubule around a given pixel and the radius defined by the line passing by both the studied pixel and the centrosome. The distribution of relative angle value as a function of the distance to the centrosome was determined and plotted with R. This distribution was also performed this time limiting the considered values to a band as shown in the figures or to a given zone like the AIZ.

### Statistical Analysis

Mann-Whitney non-parametric test was used to compare samples using GraphPad Prism software (Version 6.0). Error bars correspond to standard error mean (SEM).

**Figure 7.**
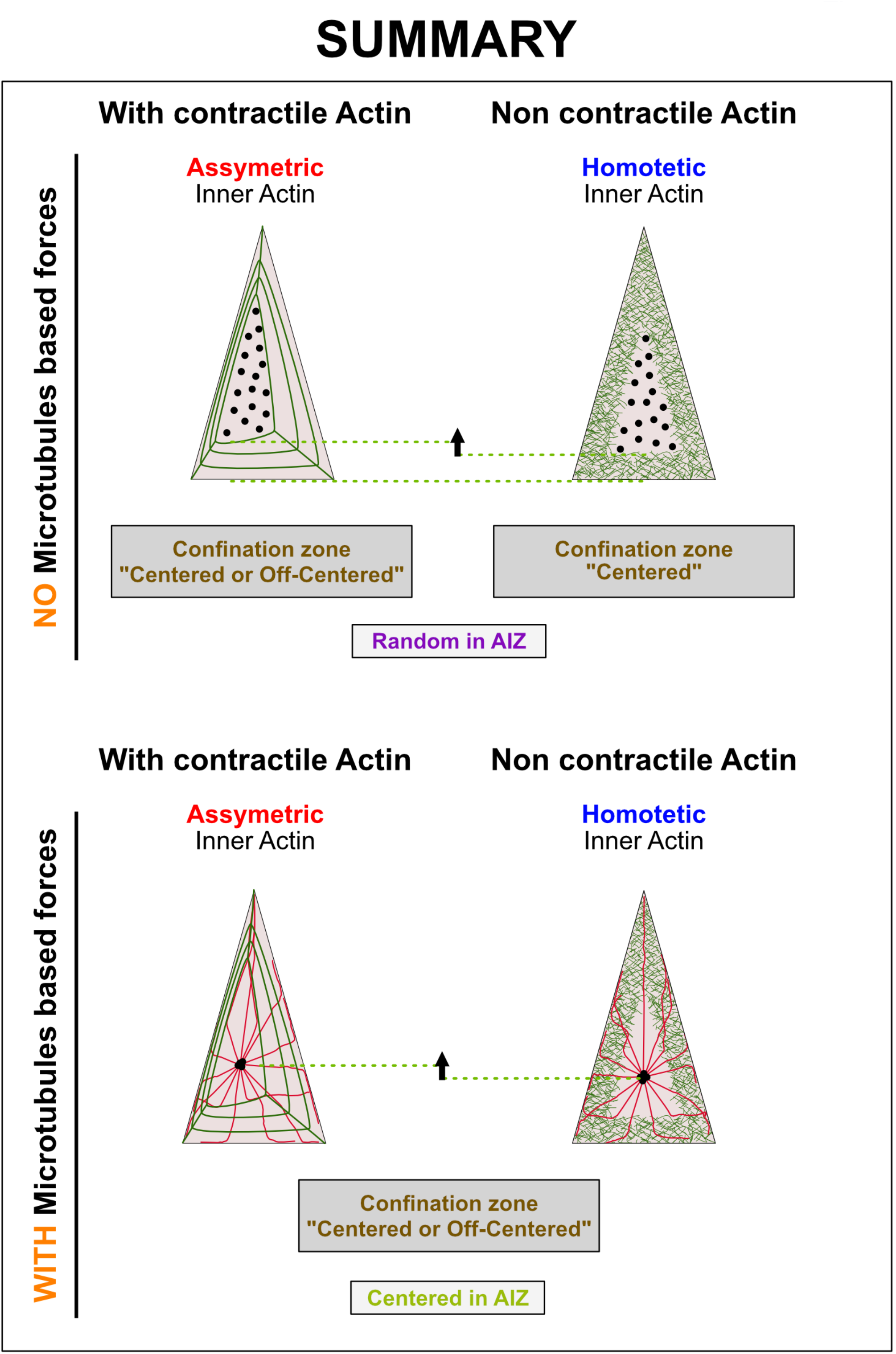
Summary. Green cables represent actin filaments. Red cables represent microtubules. Black dots represent potential centrosome positions.

## References

[1] N. Tang, W.F. Marshall, Centrosome positioning in vertebrate development, J. Cell Sci. 125 (2012) 4951–4961. https://doi.org/10.1242/jcs.038083.

[2] M. Bornens, The Centrosome in Cells and Organisms, Science. 335 (2012) 422–426. https://doi.org/10.1126/science.1209037.

[3] M. Bornens, Cell polarity: having and making sense of direction-on the evolutionary significance of the primary cilium/centrosome organ in Metazoa, Open Biol. 8 (2018). https://doi.org/10.1098/rsob.180052.

[4] M. Bornens, Organelle positioning and cell polarity, Nat. Rev. Mol. Cell Biol. 9 (2008) 874–886. https://doi.org/10.1038/nrm2524.

[5] U. Euteneuer, M. Schliwa, Mechanism of centrosome positioning during the wound response in BSC-1 cells., J. Cell Biol. 116 (1992) 1157–1166. https://doi.org/10.1083/jcb.116.5.1157.

[6] A.D. Sanchez, J.L. Feldman, Microtubule-organizing centers: from the centrosome to non-centrosomal sites, Curr. Opin. Cell Biol. 44 (2017) 93–101. https://doi.org/10.1016/j.ceb.2016.09.003.

[7] A. Müsch, Microtubule Organization and Function in Epithelial Cells, Traffic. 5 (2004) 1–9. https://doi.org/10.1111/j.1600-0854.2003.00149.x.

[8] J. Fu, I.M. Hagan, D.M. Glover, The Centrosome and Its Duplication Cycle, Cold Spring Harb. Perspect. Biol. 7 (2015) a015800. https://doi.org/10.1101/cshperspect.a015800.

[9] A. Pitaval, F. Senger, G. Letort, X. Gidrol, L. Guyon, J. Sillibourne, M. Théry, Microtubule stabilization drives 3D centrosome migration to initiate primary ciliogenesisCentrosome 3D migration during ciliogenesis, J. Cell Biol. 216 (2017) 3713–3728. https://doi.org/10.1083/jcb.201610039.

[10] D. Obino, F. Farina, O. Malbec, P.J. Sáez, M. Maurin, J. Gaillard, F. Dingli, D. Loew, A. Gautreau, M.-I. Yuseff, L. Blanchoin, M. Théry, A.-M. Lennon-Duménil, Actin nucleation at the centrosome controls lymphocyte polarity, Nat. Commun. 7 (2016) 10969. https://doi.org/10.1038/ncomms10969.

[11] M. Burute, M. Prioux, G. Blin, S. Truchet, G. Letort, Q. Tseng, T. Bessy, S. Lowell, J. Young, O. Filhol, M. Théry, Polarity Reversal by Centrosome Repositioning Primes Cell Scattering during Epithelial-to-Mesenchymal Transition, Dev. Cell. 40 (2017) 168–184. https://doi.org/10.1016/j.devcel.2016.12.004.

[12] A. Burakov, E. Nadezhdina, B. Slepchenko, V. Rodionov, Centrosome positioning in interphase cells, J. Cell Biol. 162 (2003) 963–969. https://doi.org/10.1083/jcb.200305082.

[13] J. Zhu, A. Burakov, V. Rodionov, A. Mogilner, Finding the cell center by a balance of dynein and myosin pulling and microtubule pushing: a computational study, Mol. Biol. Cell. 21 (2010) 4418–4427. https://doi.org/10.1091/mbc.E10-07-0627.

[14] L. Laan, S. Roth, M. Dogterom, End-on microtubule-dynein interactions and pulling-based positioning of microtubule organizing centers, Cell Cycle Georget. Tex. 11 (2012) 3750–3757. https://doi.org/10.4161/cc.21753.

[15] L. Laan, N. Pavin, J. Husson, G. Romet-Lemonne, M. van Duijn, M.P. López, R.D. Vale, F. Jülicher, S.L. Reck-Peterson, M. Dogterom, Cortical dynein controls microtubule dynamics to generate pulling forces that position microtubule asters, Cell. 148 (2012) 502–514. https://doi.org/10.1016/j.cell.2012.01.007.

[16] G. Letort, F. Nedelec, L. Blanchoin, M. Théry, Centrosome centering and decentering by microtubule network rearrangement, Mol. Biol. Cell. 27 (2016) 2833–2843. https://doi.org/10.1091/mbc.E16-06-0395.

[17] H. Tanimoto, J. Sallé, L. Dodin, N. Minc, Physical Forces Determining the Persistency and Centering Precision of Microtubule Asters, Nat. Phys. 14 (2018) 848–854. https://doi.org/10.1038/s41567-018-0154-4.

[18] J. Odell, V. Sikirzhytski, I. Tikhonenko, S. Cobani, A. Khodjakov, M. Koonce, Force balances between interphase centrosomes as revealed by laser ablation, Mol. Biol. Cell. 30 (2019) 1705–1715. https://doi.org/10.1091/mbc.E19-01-0034.

[19] M.P. Koonce, J. Köhler, R. Neujahr, J.-M. Schwartz, I. Tikhonenko, G. Gerisch, Dynein motor regulation stabilizes interphase microtubule arrays and determines centrosome position, EMBO J. 18 (1999) 6786–6792. https://doi.org/10.1093/emboj/18.23.6786.

[20] J. Wu, G. Misra, R.J. Russell, A.J.C. Ladd, T.P. Lele, R.B. Dickinson, Effects of dynein on microtubule mechanics and centrosome positioning, Mol. Biol. Cell. 22 (2011) 4834–4841. https://doi.org/10.1091/mbc.e11-07-0611.

[21] M. Pinot, F. Chesnel, J.Z. Kubiak, I. Arnal, F.J. Nedelec, Z. Gueroui, Effects of Confinement on the Self-Organization of Microtubules and Motors, Curr. Biol. 19 (2009) 954–960. https://doi.org/10.1016/j.cub.2009.04.027.

[22] D.A. Brito, J. Strauss, V. Magidson, I. Tikhonenko, A. Khodjakov, M.P. Koonce, Pushing Forces Drive the Comet-like Motility of Microtubule Arrays in Dictyostelium, Mol. Biol. Cell. 16 (2005) 3334–3340. https://doi.org/10.1091/mbc.e05-01-0057.

[23] J. Rosenblatt, L.P. Cramer, B. Baum, K.M. McGee, Myosin II-dependent cortical movement is required for centrosome separation and positioning during mitotic spindle assembly, Cell. 117 (2004) 361–372.

[24] C.M. Hale, W.-C. Chen, S.B. Khatau, B.R. Daniels, J.S.H. Lee, D. Wirtz, SMRT analysis of MTOC and nuclear positioning reveals the role of EB1 and LIC1 in single-cell polarization, J. Cell Sci. 124 (2011) 4267–4285. https://doi.org/10.1242/jcs.091231.

[25] M. Théry, V. Racine, M. Piel, A. Pépin, A. Dimitrov, Y. Chen, J.-B. Sibarita, M. Bornens, Anisotropy of cell adhesive microenvironment governs cell internal organization and orientation of polarity, Proc. Natl. Acad. Sci. 103 (2006) 19771–19776. https://doi.org/10.1073/pnas.0609267103.

[26] D.M. Graham, T. Andersen, L. Sharek, G. Uzer, K. Rothenberg, B.D. Hoffman, J. Rubin, M. Balland, J.E. Bear, K. Burridge, Enucleated cells reveal differential roles of the nucleus in cell migration, polarity, and mechanotransductionEnucleated cells in migration and mechanotransduction, J. Cell Biol. 217 (2018) 895–914. https://doi.org/10.1083/jcb.201706097.

[27] F.J. Ndlec, T. Surrey, A.C. Maggs, S. Leibler, Self-organization of microtubules and motors, Nature. 389 (1997) 305–308. https://doi.org/10.1038/38532.

[28] M.P.N. Juniper, M. Weiss, I. Platzman, J.P. Spatz, T. Surrey, Spherical network contraction forms microtubule asters in confinement, Soft Matter. 14 (2018) 901–909. https://doi.org/10.1039/C7SM01718A.

[29] V.I. Rodionov, G.G. Borisy, Self-centring activity of cytoplasm, Nature. 386 (1997) 170–173. https://doi.org/10.1038/386170a0.

[30] F. Pouthas, P. Girard, V. Lecaudey, T.B.N. Ly, D. Gilmour, C. Boulin, R. Pepperkok, E.G. Reynaud, In migrating cells, the Golgi complex and the position of the centrosome depend on geometrical constraints of the substratum, J. Cell Sci. 121 (2008) 2406–2414. https://doi.org/10.1242/jcs.026849.

[31] J. Zhang, Y.-L. Wang, Centrosome defines the rear of cells during mesenchymal migration, Mol. Biol. Cell. 28 (2017) 3240–3251. https://doi.org/10.1091/mbc.E17-06-0366.

[32] A.E. Rodríguez-Fraticelli, M. Auzan, M.A. Alonso, M. Bornens, F. Martín-Belmonte, Cell confinement controls centrosome positioning and lumen initiation during epithelial morphogenesis, J. Cell Biol. 198 (2012) 1011–1023. https://doi.org/10.1083/jcb.201203075.

[33] E.R. Gomes, S. Jani, G.G. Gundersen, Nuclear movement regulated by Cdc42, MRCK, myosin, and actin flow establishes MTOC polarization in migrating cells, Cell. 121 (2005) 451–463. https://doi.org/10.1016/j.cell.2005.02.022.

[34] I. Dupin, E. Camand, S. Etienne-Manneville, Classical cadherins control nucleus and centrosome position and cell polarity, J. Cell Biol. 185 (2009) 779–786. https://doi.org/10.1083/jcb.200812034.

[35] M. Almonacid, W.W. Ahmed, M. Bussonnier, P. Mailly, T. Betz, R. Voituriez, N.S. Gov, M.-H. Verlhac, Active diffusion positions the nucleus in mouse oocytes, Nat. Cell Biol. 17 (2015) 470–479. https://doi.org/10.1038/ncb3131.

[36] I. Dupin, Y. Sakamoto, S. Etienne-Manneville, Cytoplasmic intermediate filaments mediate actin-driven positioning of the nucleus, J. Cell Sci. 124 (2011) 865–872. https://doi.org/10.1242/jcs.076356.

[37] I. Dupin, S. Etienne-Manneville, Nuclear positioning: Mechanisms and functions, Int. J. Biochem. Cell Biol. 43 (2011) 1698–1707. https://doi.org/10.1016/j.biocel.2011.09.004.

[38] A.V. Burakov, E.S. Nadezhdina, Association of nucleus and centrosome: magnet or velcro?, Cell Biol. Int. 37 (2013) 95–104. https://doi.org/10.1002/cbin.10016.

[39] M. Bornens, Is the centriole bound to the nuclear membrane?, Nature. 270 (1977) 80–82. https://doi.org/10.1038/270080a0.

[40] G. Salpingidou, A. Smertenko, I. Hausmanowa-Petrucewicz, P.J. Hussey, C.J. Hutchison, A novel role for the nuclear membrane protein emerin in association of the centrosome to the outer nuclear membrane, J. Cell Biol. 178 (2007) 897–904. https://doi.org/10.1083/jcb.200702026.

[41] B.M. Maro B, The centriole–nucleus association: effects of cytochalasin B and nocodazole, Biol. Cell. (n.d.) 287–90.

[42] J.W. Shay, K.R. Porter, D.M. Prescott, The surface morphology and fine structure of CHO (Chinese hamster ovary) cells following enucleation, Proc. Natl. Acad. Sci. U. S. A. 71 (1974) 3059–3063. https://doi.org/10.1073/pnas.71.8.3059.

[43] E. Karsenti, S. Kobayashi, T. Mitchison, M. Kirschner, Role of the centrosome in organizing the interphase microtubule array: properties of cytoplasts containing or lacking centrosomes, J. Cell Biol. 98 (1984) 1763–1776. https://doi.org/10.1083/jcb.98.5.1763.

[44] X. Zhang, K. Lei, X. Yuan, X. Wu, Y. Zhuang, T. Xu, R. Xu, M. Han, SUN1/2 and Syne/Nesprin-1/2 Complexes Connect Centrosome to the Nucleus during Neurogenesis and Neuronal Migration in Mice, Neuron. 64 (2009) 173–187. https://doi.org/10.1016/j.neuron.2009.08.018.

[45] C.J. Malone, L. Misner, N.L. Bot, M.-C. Tsai, J.M. Campbell, J. Ahringer, J.G. White, The C. elegans Hook Protein, ZYG-12, Mediates the Essential Attachment between the Centrosome and Nucleus, Cell. 115 (2003) 825–836. https://doi.org/10.1016/S0092-8674(03)00985-1.

[46] D.A. Starr, H.N. Fridolfsson, Interactions Between Nuclei and the Cytoskeleton Are Mediated by SUN-KASH Nuclear-Envelope Bridges, Annu. Rev. Cell Dev. Biol. 26 (2010) 421–444. https://doi.org/10.1146/annurev-cellbio-100109-104037.

[47] H. Umeshima, T. Hirano, M. Kengaku, Microtubule-based nuclear movement occurs independently of centrosome positioning in migrating neurons, Proc. Natl. Acad. Sci. U. S. A. 104 (2007) 16182–16187. https://doi.org/10.1073/pnas.0708047104.

[48] P.J. Strzyz, H.O. Lee, J. Sidhaye, I.P. Weber, L.C. Leung, C. Norden, Interkinetic nuclear migration is centrosome independent and ensures apical cell division to maintain tissue integrity, Dev. Cell. 32 (2015) 203–219. https://doi.org/10.1016/j.devcel.2014.12.001.

[49] R.A. Battaglia, S. Delic, H. Herrmann, N.T. Snider, Vimentin on the move: new developments in cell migration, F1000Research. 7 (2018). https://doi.org/10.12688/f1000research.15967.1.

[50] B.T. Helfand, M.G. Mendez, S.N.P. Murthy, D.K. Shumaker, B. Grin, S. Mahammad, U. Aebi, T. Wedig, Y.I. Wu, K.M. Hahn, M. Inagaki, H. Herrmann, R.D. Goldman, Vimentin organization modulates the formation of lamellipodia, Mol. Biol. Cell. 22 (2011) 1274–1289. https://doi.org/10.1091/mbc.E10-08-0699.

[51] M.G. Mendez, S.-I. Kojima, R.D. Goldman, Vimentin induces changes in cell shape, motility, and adhesion during the epithelial to mesenchymal transition, FASEB J. 24 (2010) 1838–1851. https://doi.org/10.1096/fj.09-151639.

[52] M. Piel, P. Meyer, A. Khodjakov, C.L. Rieder, M. Bornens, The respective contributions of the mother and daughter centrioles to centrosome activity and behavior in vertebrate cells, J. Cell Biol. 149 (2000) 317–330. https://doi.org/10.1083/jcb.149.2.317.

[53] Triangle center, Wikipedia. (2019). https://en.wikipedia.org/w/index.php?title=Triangle_center&oldid=931987767 (accessed January 7, 2020).

[54] M. Théry, A. Pépin, E. Dressaire, Y. Chen, M. Bornens, Cell distribution of stress fibres in response to the geometry of the adhesive environment, Cell Motil. 63 (2006) 341–355. https://doi.org/10.1002/cm.20126.

[55] T. Chen, A. Callan-Jones, E. Fedorov, A. Ravasio, A. Brugués, H.T. Ong, Y. Toyama, B.C. Low, X. Trepat, T. Shemesh, R. Voituriez, B. Ladoux, Large-scale curvature sensing by directional actin flow drives cellular migration mode switching, Nat. Phys. 15 (2019) 393–402. https://doi.org/10.1038/s41567-018-0383-6.

[56] S.W. Grill, A.A. Hyman, Spindle positioning by cortical pulling forces, Dev. Cell. 8 (2005) 461–465. https://doi.org/10.1016/j.devcel.2005.03.014.

[57] B.P. Liu, M. Chrzanowska-Wodnicka, K. Burridge, Microtubule depolymerization induces stress fibers, focal adhesions, and DNA synthesis via the GTP-binding protein Rho, Cell Adhes. Commun. 5 (1998) 249–255. https://doi.org/10.3109/15419069809040295.

[58] A. De Simone, F. Nédélec, P. Gönczy, Dynein Transmits Polarized Actomyosin Cortical Flows to Promote Centrosome Separation, Cell Rep. 14 (2016) 2250–2262. https://doi.org/10.1016/j.celrep.2016.01.077.

[59] Y.K. Wu, H. Umeshima, J. Kurisu, M. Kengaku, Nesprins and opposing microtubule motors generate a point force that drives directional nuclear motion in migrating neurons, Development. 145 (2018). https://doi.org/10.1242/dev.158782.

[60] G.G. Luxton, G.G. Gundersen, Orientation and function of the nuclear–centrosomal axis during cell migration, Curr. Opin. Cell Biol. 23 (2011) 579–588. https://doi.org/10.1016/j.ceb.2011.08.001.

[61] M. Dogterom, G.H. Koenderink, Actin–microtubule crosstalk in cell biology, Nat. Rev. Mol. Cell Biol. 20 (2019) 38–54. https://doi.org/10.1038/s41580-018-0067-1.

[62] S.L. Gupton, W.C. Salmon, C.M. Waterman-Storer, Converging populations of f-actin promote breakage of associated microtubules to spatially regulate microtubule turnover in migrating cells, Curr. Biol. CB. 12 (2002) 1891–1899. https://doi.org/10.1016/s0960-9822(02)01276-9.

[63] A. Colin, P. Singaravelu, M. Théry, L. Blanchoin, Z. Gueroui, Actin-Network Architecture Regulates Microtubule Dynamics, Curr. Biol. CB. 28 (2018) 2647–2656.e4. https://doi.org/10.1016/j.cub.2018.06.028.

[64] R.D. Vale, Y.Y. Toyoshima, Rotation and translocation of microtubules in vitro induced by dyneins from Tetrahymena cilia, Cell. 52 (1988) 459–469. https://doi.org/10.1016/S0092-8674(88)80038-2.

[65] P. Gönczy, S. Pichler, M. Kirkham, A.A. Hyman, Cytoplasmic dynein is required for distinct aspects of MTOC positioning, including centrosome separation, in the one cell stage Caenorhabditis elegans embryo, J. Cell Biol. 147 (1999) 135–150. https://doi.org/10.1083/jcb.147.1.135.

[66] K. Kimura, A. Kimura, A novel mechanism of microtubule length-dependent force to pull centrosomes toward the cell center, Bioarchitecture. 1 (2011) 74–79. https://doi.org/10.4161/bioa.1.2.15549.

[67] I. Kimura, D. Inoue, T. Maeda, T. Hara, A. Ichimura, S. Miyauchi, M. Kobayashi, A. Hirasawa, G. Tsujimoto, Short-chain fatty acids and ketones directly regulate sympathetic nervous system via G protein-coupled receptor 41 (GPR41), Proc. Natl. Acad. Sci. U. S. A. 108 (2011) 8030–8035. https://doi.org/10.1073/pnas.1016088108.

