## Supplementary figures and images for "Acto-myosin network geometry defines centrosome position"

### Supplementary Figures

#### Figure S1

A

Anti-Vimentin

Anti-Vimentin

Cytoplasts RPE1

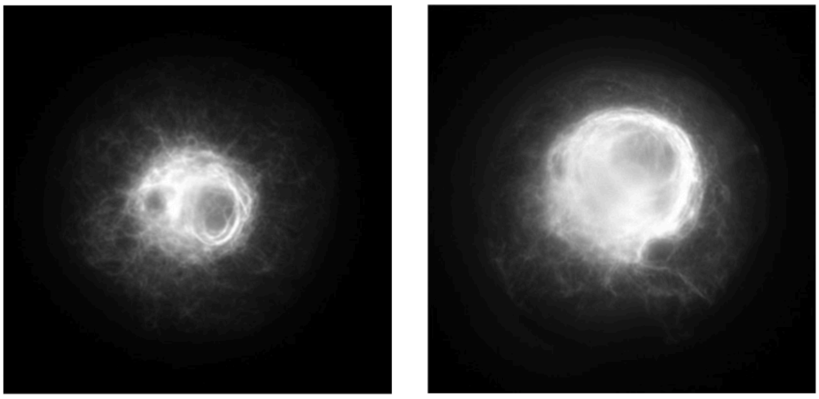

B

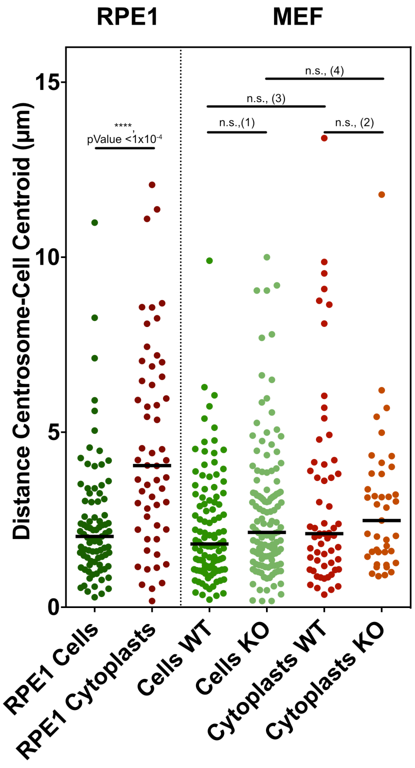


*(1), pValue = 0.0638 ; (2), pValue = 0.7277 ; (3), pValue = 0.1413 ; (4), pValue = 0.5659*

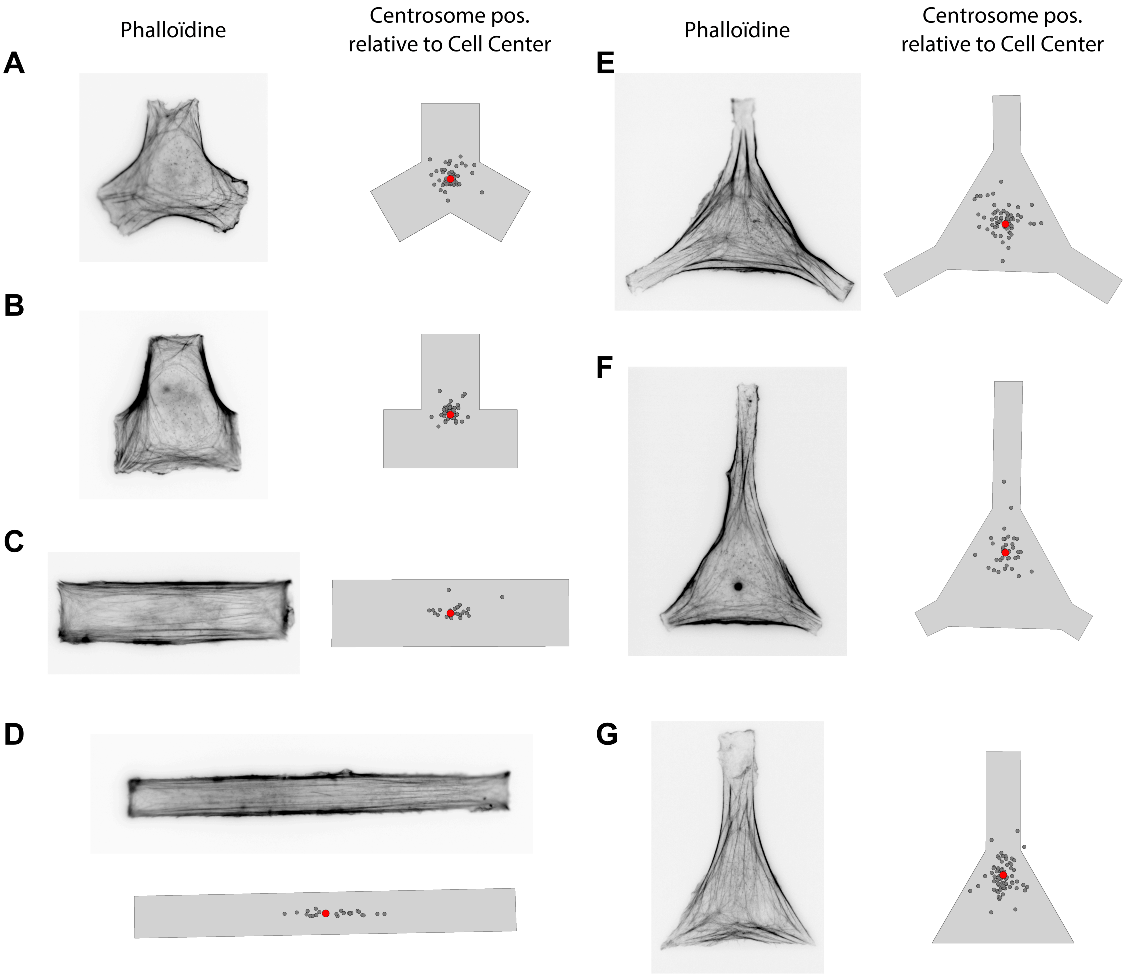

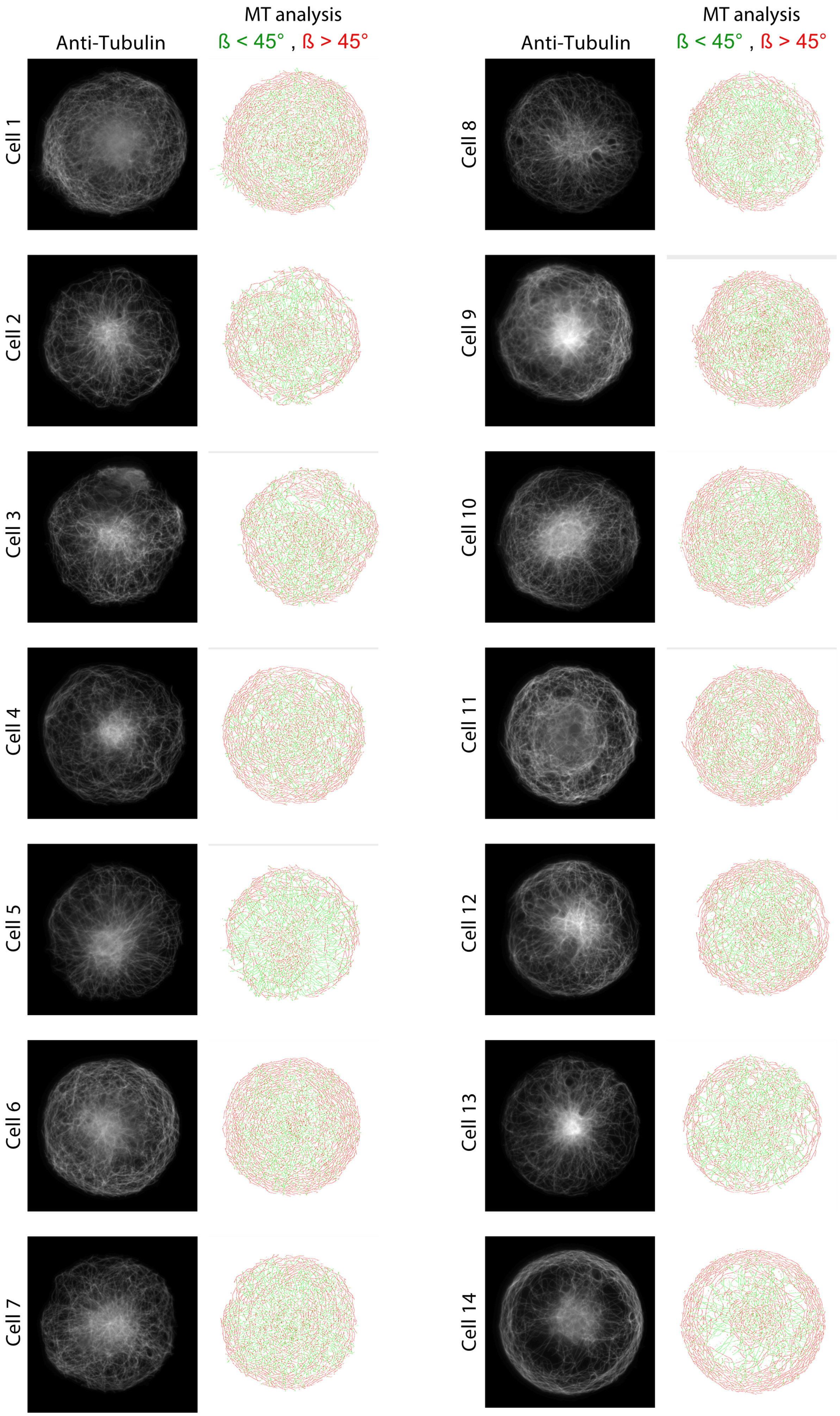

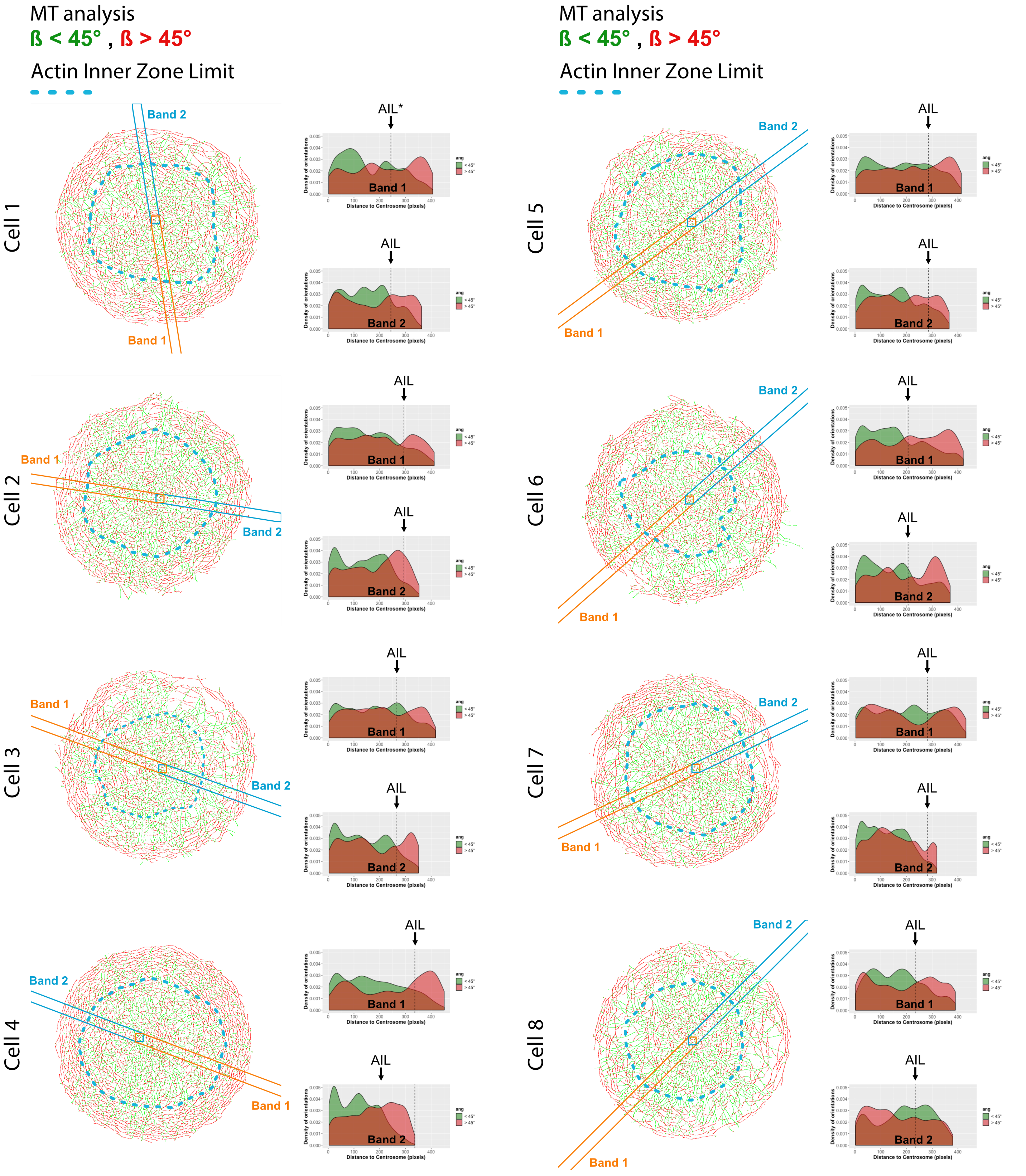

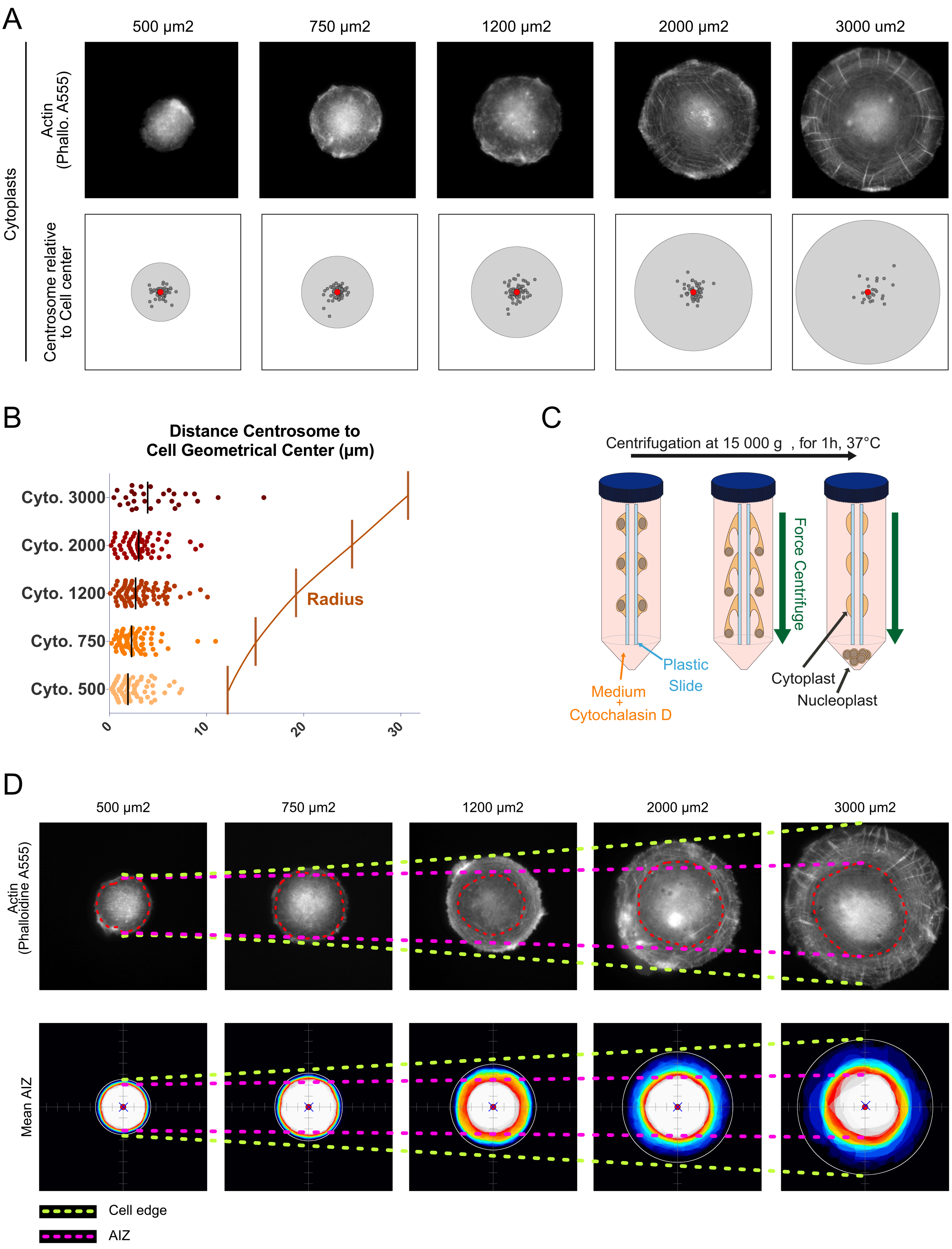

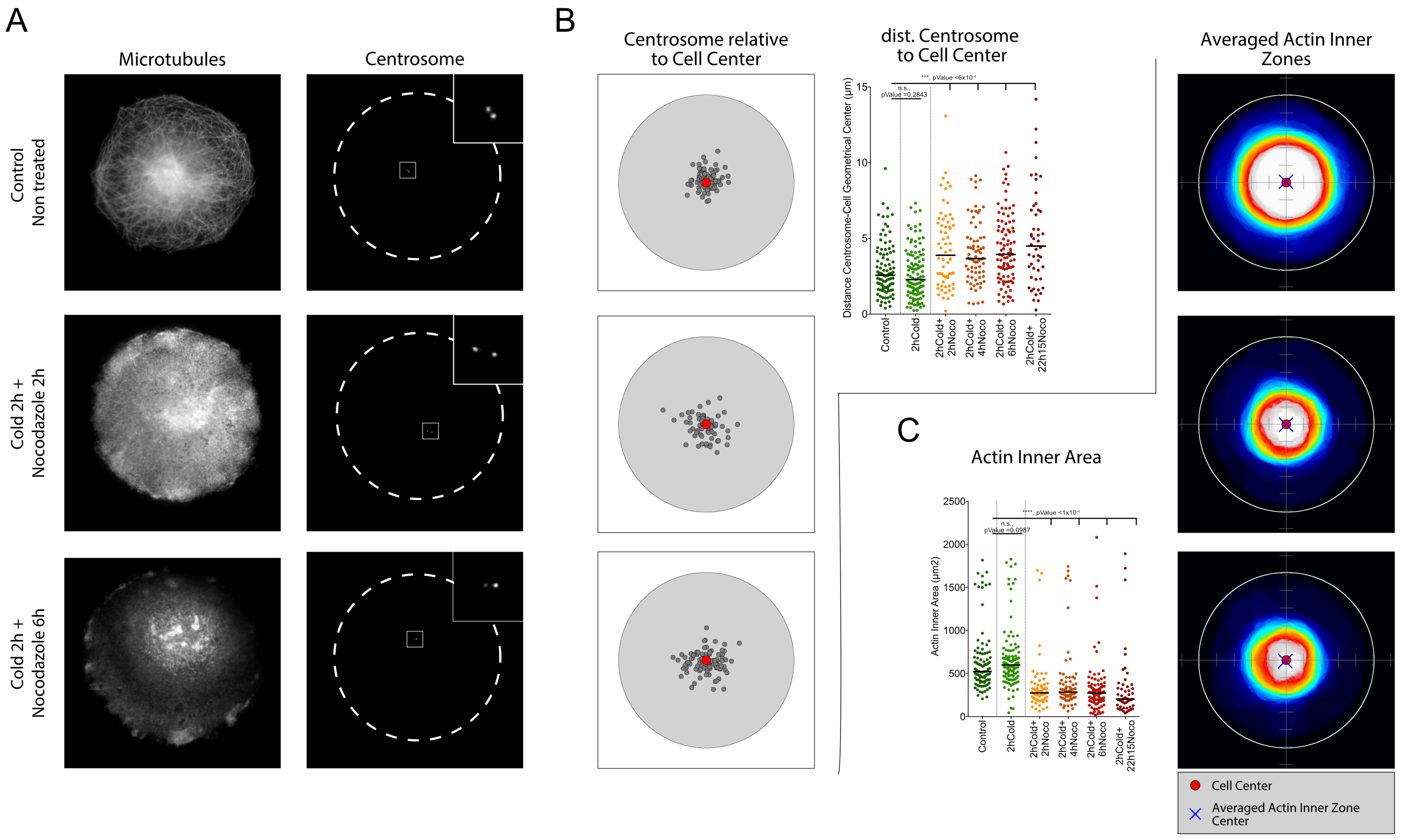
