## Supplementary Legends for "Acto-myosin network geometry defines centrosome position"

**Legends for supplementary figures**

Figure S1. **Vimentin filaments form circular structures upon enucleation.**
**A.** Cytoplasts made with RPE1 cells present Vimentin cage-like structures, which most likely explain differences in centrosome position distribution between cells and cytoplasts.
**B.** Quantification of centrosome position for three cell lines plated in 2000µm2 disks: RPE1, MEF WT or MEF KO for Vimentin. RPE1 cells (which contain high levels of Vimentin) and cytoplasts present very different distribution of centrosome positioning. On the other hand, MEF WT and KO for Vimentin and their respective cytoplasts present very similar distribution, also similar to RPE1 cells. The lack of difference between MEF WT and its corresponding cytoplasts may be due to the natural low levels of Vimentin present in MEF WT cells.

Figure S2. **The centrosome is close to the cell center in most shapes.**
Cytoplasts from MEF KO Vimentin cells were seeded on 2000µm2 micropatterns with a variety of shapes. They were stained for centrosome and actin, and centrosome positioning was assed and plotted relative to the cell geometrical center. Overlapping images were taken and stitched for D,F and G before image processing.

Figure S3. **Microtubule network systematic analysis**
Several examples of the analysis of microtubule network orientation are presented. Cytoplasts spread on 2000µm2 disks were fixed and stained for microtubules and centrosome. Image stacks were taken and analyzed using D-FiNS, using the centrosome coordinates as the center for the calculation of the relative orientation. Best plane of microtubules in shown on the left columns and the orientation map is shown on the right columns.

Figure S4. **Microtubule network systematic analysis crossed with AIZ analysis**
Several examples of the analysis of microtubule network orientation are presented. Cytoplasts spread on 2000µm2 disks were fixed and stained for microtubules, actin and centrosome. AIZ was determined for each cell and microtubule network orientation was performed within two bands, starting from the centrosome and defined by the diameter passing by both the cell centroid and the centrosome. Radial and Tangential orientation density as a function of the distance to the centrosome were plotted using R software. AIL stands for Actin Inner Limit.

Figure S5. **Centrosome positioning does not depend on spreading size in disks.**
Cytoplasts were platted on disks of different sizes: 500, 750, 1200, 2000, 3000µm2.
**A.** 83% of the centrosomes were found in a 5µm wide region around the geometrical center of the cell.
**B.** Compared to radius increase, the distance of the centrosome to the cell center remains stationary.
**C.** Scheme summarizing cytoplasts production protocole.
**D.** Compared to the increase of the cell diameter, the diameter of the average actin inner zone remains rather constant for all pattern sizes.

Figure S6. **Centrosome dispersion due to absence of microtubules is limited by centripetal contractility increase in disks.**
**A.** Efficient removal of microtubules is achieved by incubation on ice for 2 hours followed by warming up in the presence of 10µM Nocodazole and incubation for different lapses of time.

**B.** Centrosome distance to cell center is higher in Ice-Nocodazole conditions but is not randomized all over the cell as expected.
**C.** AIZs are smaller and centered in Ice-Nocodazole conditions, probably explaining the limited dispersion of centrosomes.
Red dot represents cell geometrical center or the AIC as indicated by the graph title. In **C**, the red dot represents the cell geometrical center and the blue cross represents the averaged relative AIC. The grey dots are the centrosomes.
